# Dysregulated Expression of Circadian Genes in Lymphoblastoid Cells of Patients with ASD

**DOI:** 10.1101/2022.10.03.510736

**Authors:** Hui Ding, Shuzhen Kuang, Chin-Fu Chen

**Affiliations:** San Diego, CA, USA; San Francisco, CA, USA; Sinceris Consulting, Clemson, SC, 29631, USA

**Keywords:** Autism Spectrum Disorder (ASD), circadian rhythm, circadian gene expression, co-expression network, lymphoblastoid cells

## Abstract

Patients with Autism Spectrum Disorder (ASD) often exhibit disturbances in sleep, metabolism, and immune system. The molecular mechanisms for these clinical features in ASD are currently unknown. We demonstrated that circadian genes in the cells of patients with ASD often are dysregulated compared to controls. The dysregulation of circadian genes was reflected in two different ways: (1) abnormal levels of expression; and (2) a change of gene-gene association pattern in the co-expression network. We also observed a link between abnormal expression of circadian genes in lymphoblastoid cells with sleep phenotypes in patients with ASD. Our results suggest that circadian genes and circadian rhythms might play critical roles in the pathogenesis of ASD.

## Introduction

Autism Spectrum Disorder (ASD) is a neurodevelopmental disorder characterized by deficits in communication and social interaction. ASD is highly heritable (Berg and Geschwind, 2012; Hallmayer et al., 2011; Ronald and Hoekstra, 2011; Sandin et al., 2014) but its genetic architecture is complex. First, more than 100 ASD risk genes have been proposed (Betancur, 2011; Devlin and Scherer, 2012). Further, only 20-25% of patients with AD can be diagnosed with a genetic defect, the cause for the rest of 75-80% is unknown (Miles, 2011). Despite the genetic complexity of ASD, several clinical features are common in patients with ASD: (1) Sleep disorders (in∼44-83% of cases) (Johnson and Malow, 2008); (2) Immune system related symptoms and inflammatory disorders (Estes and McAllister, 2015; Hsiao, 2013); (3) Metabolic abnormalities (Frye, 2015).

We noticed that these co-appearing phenotypes of ASD resemble some features in patients with disorders in circadian rhythm. Recent studies of other human diseases demonstrated that disturbed circadian rhythms are associated with neuropsychiatric disorders (Lamont et al., 2010; Liu and Chung, 2015), diabetes (Nagorny and Lyssenko, 2012), obesity (Froy, 2010), metabolic syndrome (Maury et al., 2010), and immune dysfunction (Curtis et al., 2014; Swanson et al., 2011). It is likely that patients with ASD having clinical features in sleep, immune system, and metabolism may contain defects in the molecular pathways that control circadian rhythm.

The circadian system in humans is organized into molecular oscillators, also known as circadian molecular clocks, in the brain and throughout the body. The master clock is the suprachiasmatic nucleus (SCN) in the hypothalamus. External stimuli such as light or food, can also modulate or reset the circadian clock. How the SCN coordinates and synchronizes circadian rhythm with peripheral tissues is not entirely clear at present (Saini et al., 2011). However, the circadian molecular clocks, are cell-autonomous in the sense that they can function without external inputs such as light. Different cultured human cells, including fibroblasts (Brown et al., 2005), mesenchymal- and adipose-derived stem cells (Huang et al., 2009), red blood cells (O’Neill and Reddy, 2011), islet cells (Pulimeno et al., 2013), and hair follicle (Hardman et al., 2015), all have been shown to display independent circadian rhythms without changes in light or temperature conditions.

To gain insights on the functional roles of circadian rhythm in the pathogenesis of ASD, we thus investigated whether there are molecular defects of circadian system in patients with ASD. More specifically, we asked the questions: what are the levels of expressions of circadian genes in lymphoblastoid cells of patients with ASD, compared to controls? Are the expression levels among circadian genes correlated with each other in lymphoblastoid cells of patients with ASD, compared to controls?

## Results

### Gene expression levels of selected circadian genes in the lymphoblastoid cells of normal controls

It is currently unknown whether circadian genes express in the human lymphoblastoid cells. We hence selected and assayed the expression levels of ten circadian genes. The genes are: BMAL1, CLOCK, PER1, PER2, PER3, CRY1, CRY2, RORA, REV-ERBα, and NPAS2. These genes are known to play key roles in initiating and maintaining the molecular oscillator in other cell types.

Our results showed that the levels of mRNA expression for most of the selected circadian genes were within the range of other non-circadian genes (**Figure 1**). Because all the selected genes also function as transcription factors in other cell types, we suspected that circadian genes were likely to mediate gene regulation in human lymphoblastoid cells.

**Figure 1.**
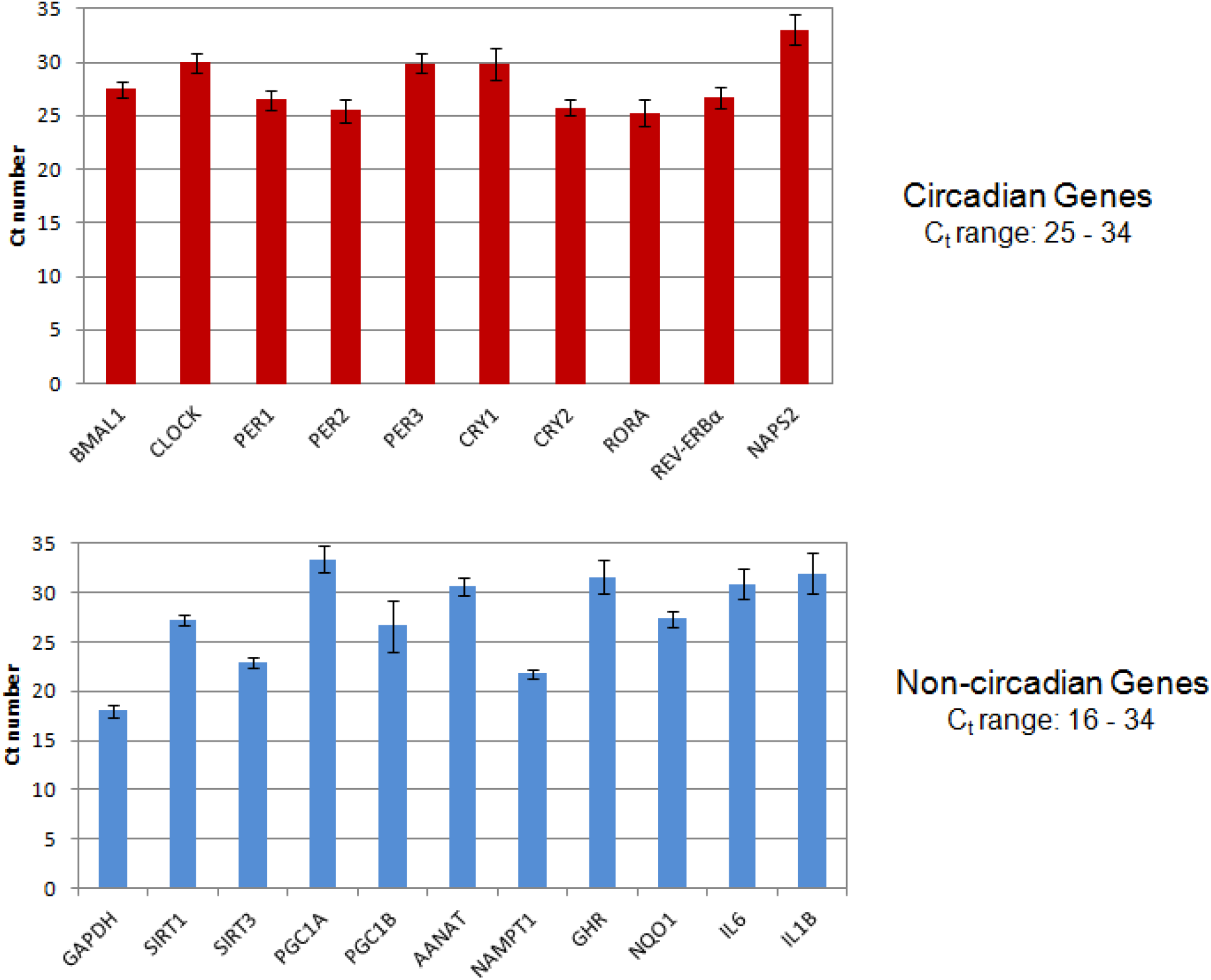
Comparison of C_t_ numbers of qPCR for 10 selected circadian genes and 11 non-circadian genes. The bar height represents mean (of three technical replicates) ± standard deviation (small vertical bars). Note: C_t_ (threshold cycle) is a relative measure of the concentration of DNA in the PCR reaction. Lower numbers indicate higher copy numbers of DNA in the reaction.

### Abnormal gene expression in the lymphoblastoid cells of patients with ASD

We next compared gene expressions of the selected circadian genes in the lymphoblastoid cells of patients with ASD with those of controls. Our results suggest that many patients with ASD display dramatic increased or decreased expressions of multiple circadian genes (**Table 1**). On average, a patient with ASD had about 1.5 out of the 10 circadian genes being ‘abnormal’, as defined by a change equal to or greater than 2 standard deviations from the mean value. In contrast, a normal person on average will have only about 0.4 out of the 10 circadian genes that would be abnormal. In other words, the chance of having an abnormal level of circadian gene expression for a patient with ASD is almost four times more than the controls. The odds ratio of a patient with ASD having an abnormal circadian gene is 3.85 (95% confidence levels 2.13 to 6.96).

**Table 1.**
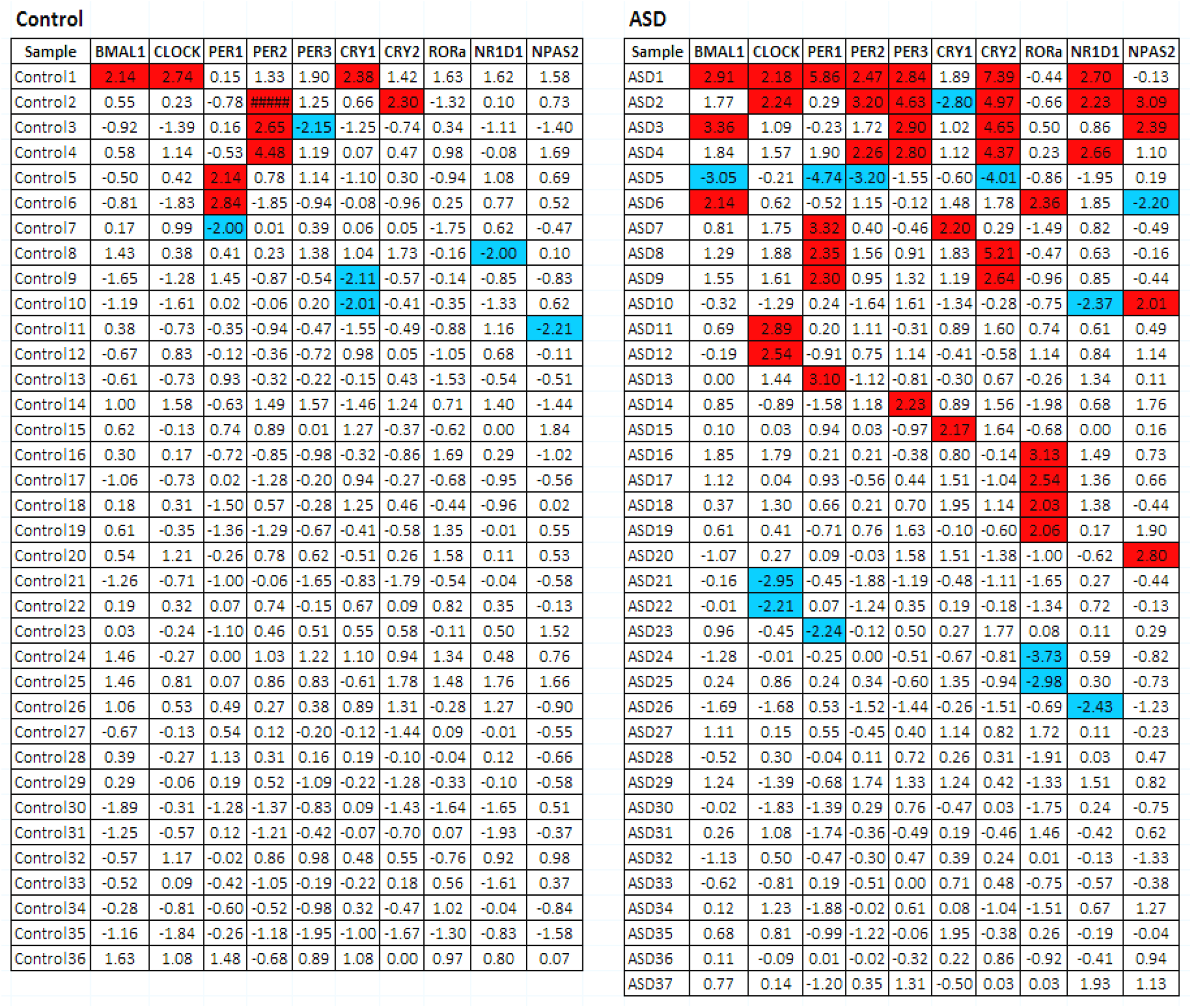
Z-scores of qPCR results of 10 selected circadian genes in the lymphoblastoid cells of controls (left panel) versus that of patients with ASD (right panel). Red background color denotes Z score > 2 and blue color indicate Z score < −2. Note: A Z-score of 2 represents ∼2.3% of the area on the right tail of a Z distribution, while a Z-score of −2 denotes ∼2.3% of the area on the right tail of the distribution.

Overall, more than seventy percent of patients with ASD (26 out of 37) have one or more genes out of the normal range. In contrast, only about thirty percent of controls (11 out of 36) have one or more genes out of the normal range. Furthermore, 6 out of the 38 patients with ASD have more than 3 circadian genes with abnormal expression (∼16.2%), while only one person out the 36 controls has 3 circadian genes with abnormal expression (∼2.8%).

### Abnormal gene co-expression network in the lymphoblastoid cells of patients with ASD

Genes do not function in isolation. Genes can form interconnected networks (associations) at the transcriptional levels. The functional roles of genes thus are determined by the context of their relationships with other genes within the network. We believe that co-expression networks, as often described as guilt-by-association, can elucidate the biological context of gene regulation. We computed the degree of association between each gene of interest via pair-wise Spearman correlation coefficient which is less sensitive to the outliers. The minimal strength of association (measured by the correlation coefficient) to be depicted in the network was arbitrarily chosen at 0.6 (*P* < 0.001).

Inspection of the co-expression network of the 10 circadian genes for the control cells revealed that one prominent feature was the four inter-locked loops of BMAL1-CLOCK-CRY2-PER3 (**Figure 2**, left panel). For other part of the network, PER2 had a single connection with CRY2, while PER1, REV-ERBα, RORA, and NAPS2, and CRY1 all were singletons without strong connections to any other members of circadian genes (**Figure 2**, left panel).

**Figure 2.**
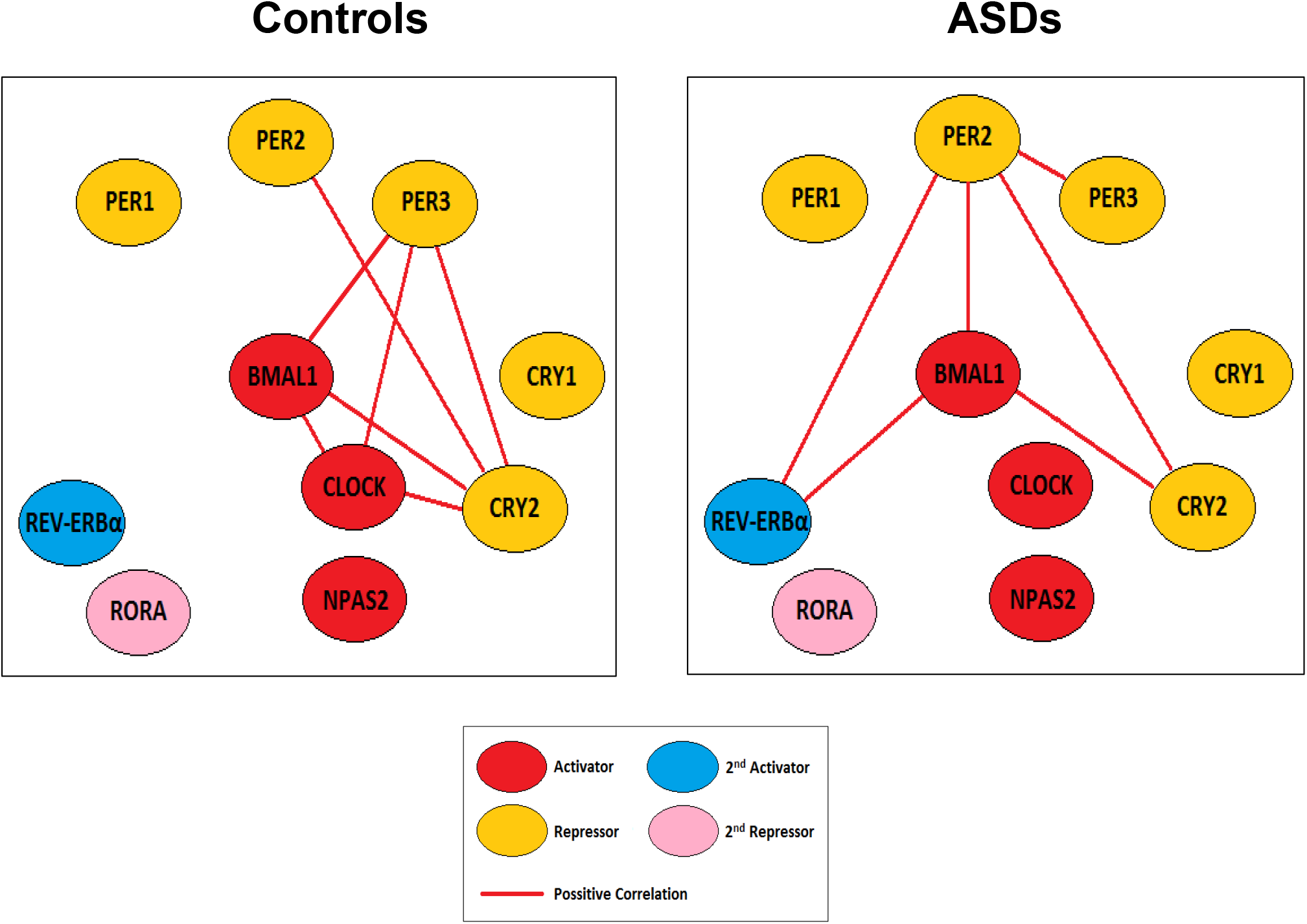
Co-expression networks of 10 circadian genes in the lymphoblastoid cells of controls versus that of patients with ASD. The edges denote Spearman coefficient *r* ≥0.6 (*P* < 0.0001) between the nodes. The color-coded roles of different circadian genes, such as activators and repressors, are inferred from other cell types.

The co-expression network of the circadian genes for the ASD cells is very different from that of the controls. There were losses of connections as well as gains of connections between the 10 circadian genes, compared to those of the control. The co-expression network of the circadian genes in the ASD cells was dominated with a different four-partner BMAL1-CRY2-PER2-REV-ERBα loops (**Figure 2**, right panel). One striking feature was the complete loss of connection for CLOCK gene to any other genes in the ASD network. Another notable characteristics was the newly gained connections of REV-ERBα with other genes in the ASD network. PER2 also had a dramatic change: it was connected with multiple partners. On the other hand, PER3 was no longer an important node as in the control’s network and only had a single connection to PER2 in the ASD network. Most of the original singletons in the control’s network remain, except that REV-ERBα was no longer a singleton in the ASD’s network (**Figure 2**, right panel).

### Circadian gene expression and sleep disturbance

Disrupted circadian rhythms can lead to sleep disorders. We examined the clinical records of our ASD cohort and found that about sixty percent of (22/37) of the patients were reported with sleep problems (**Table 2**). We next asked the question if the abnormal expression levels of the cultured lymphoblastoid cells are linked to sleep phenotypes. As depicted in **Figure 3**, a majority of the patients with sleep problems could be separated from the patients with normal sleep pattern, based on data of circadian gene expressions in the lymphoblastoid cells. The results thus suggest that sleep phenotypes are correlated with the circadian gene expressions in the lymphoblastoid cells of patients with ASD.

**Table 2.** Patients with ASD with sleep problems and the circadian gene expressions (as Z-Scores).

| Sample | BMAL1 | CLOCK | PER1 | PER2 | PER3 | CRY1 | CRY2 | RORa | NR1D1 | NPAS2 | Sleep problems |
| --- | --- | --- | --- | --- | --- | --- | --- | --- | --- | --- | --- |
| ASD1 | 2.91 | 2.18 | 5.86 | 2.47 | 2.84 | 1.89 | 7.39 | -0.44 | 2.70 | -0.13 | Yes |
| ASD2 | 1.77 | 2.24 | 0.29 | 3.20 | 4.63 | -2.80 | 4.97 | -0.66 | 2.23 | 3.09 | Yes |
| ASD3 | 3.36 | 1.09 | -0.23 | 1.72 | 2.90 | 1.02 | 4.65 | 0.50 | 0.86 | 2.39 | Yes |
| ASD5 | -3.05 | -0.21 | -4.74 | -3.20 | -1.55 | -0.60 | -4.01 | -0.86 | -1.95 | 0.19 | Yes |
| ASD6 | 2.14 | 0.62 | -0.52 | 1.15 | -0.12 | 1.48 | 1.78 | 2.36 | 1.85 | -2.20 | Yes |
| ASD8 | 1.29 | 1.88 | 2.35 | 1.56 | 0.91 | 1.83 | 5.21 | -0.47 | 0.63 | -0.16 | Yes |
| ASD10 | -0.32 | -1.29 | 0.24 | -1.64 | 1.61 | -1.34 | -0.28 | -0.75 | -2.37 | 2.01 | Yes |
| ASD11 | 0.69 | 2.89 | 0.20 | 1.11 | -0.31 | 0.89 | 1.60 | 0.74 | 0.61 | 0.49 | Yes |
| ASD13 | 0.00 | 1.44 | 3.10 | -1.12 | -0.81 | -0.30 | 0.67 | -0.26 | 1.34 | 0.11 | Yes |
| ASD14 | 0.85 | -0.89 | -1.58 | 1.18 | 2.23 | 0.89 | 1.56 | -1.98 | 0.68 | 1.76 | Yes |
| ASD17 | 1.12 | 0.04 | 0.93 | -0.56 | 0.44 | 1.51 | -1.04 | 2.54 | 1.36 | 0.66 | Yes |
| ASD18 | 0.37 | 1.30 | 0.66 | 0.21 | 0.70 | 1.95 | 1.14 | 2.03 | 1.38 | -0.44 | Yes |
| ASD19 | 0.61 | 0.41 | -0.71 | 0.76 | 1.63 | -0.10 | -0.60 | 2.06 | 0.17 | 1.90 | Yes |
| ASD21 | -0.16 | -2.95 | -0.45 | -1.88 | -1.19 | -0.48 | -1.11 | -1.65 | 0.27 | -0.44 | Yes |
| ASD22 | -0.01 | -2.21 | 0.07 | -1.24 | 0.35 | 0.19 | -0.18 | -1.34 | 0.72 | -0.13 | Yes |
| ASD23 | 0.96 | -0.45 | -2.24 | -0.12 | 0.50 | 0.27 | 1.77 | 0.08 | 0.11 | 0.29 | Yes |
| ASD26 | -1.69 | -1.68 | 0.53 | -1.52 | -1.44 | -0.26 | -1.51 | -0.69 | -2.43 | -1.23 | Yes |
| ASD27 | 1.11 | 0.15 | 0.55 | -0.45 | 0.40 | 1.14 | 0.82 | 1.72 | 0.11 | -0.23 | Yes |
| ASD29 | 1.24 | -1.39 | -0.68 | 1.74 | 1.33 | 1.24 | 0.42 | -1.33 | 1.51 | 0.82 | Yes |
| ASD32 | -1.13 | 0.50 | -0.47 | -0.30 | 0.47 | 0.39 | 0.24 | 0.01 | -0.13 | -1.33 | Yes |
| ASD33 | -0.62 | -0.81 | 0.19 | -0.51 | 0.00 | 0.71 | 0.48 | -0.75 | -0.57 | -0.38 | Yes |
| ASD35 | 0.68 | 0.81 | -0.99 | -1.22 | -0.06 | 1.95 | -0.38 | 0.26 | -0.19 | -0.04 | Yes |
| ASD9 | 1.55 | 1.61 | 2.30 | 0.95 | 1.32 | 1.19 | 2.64 | -0.96 | 0.85 | -0.44 | No |
| ASD12 | -0.19 | 2.54 | -0.91 | 0.75 | 1.14 | -0.41 | -0.58 | 1.14 | 0.84 | 1.14 | No |
| ASD15 | 0.10 | 0.03 | 0.94 | 0.03 | -0.97 | 2.17 | 1.64 | -0.68 | 0.00 | 0.16 | No |
| ASD24 | -1.28 | -0.01 | -0.25 | 0.00 | -0.51 | -0.67 | -0.81 | -3.73 | 0.59 | -0.82 | No |
| ASD25 | 0.24 | 0.86 | 0.24 | 0.34 | -0.60 | 1.35 | -0.94 | -2.98 | 0.30 | -0.73 | No |
| ASD28 | -0.52 | 0.30 | -0.04 | 0.11 | 0.72 | 0.26 | 0.31 | -1.91 | 0.03 | 0.47 | No |
| ASD30 | -0.02 | -1.83 | -1.39 | 0.29 | 0.76 | -0.47 | 0.03 | -1.75 | 0.24 | -0.75 | No |
| ASD31 | 0.26 | 1.08 | -1.74 | -0.36 | -0.49 | 0.19 | -0.46 | 1.46 | -0.42 | 0.62 | No |
| ASD34 | 0.12 | 1.23 | -1.88 | -0.02 | 0.61 | 0.08 | -1.04 | -1.51 | 0.67 | 1.27 | No |
| ASD36 | 0.11 | -0.09 | 0.01 | -0.02 | -0.32 | 0.22 | 0.86 | -0.92 | -0.41 | 0.94 | No |
| ASD4 | 1.84 | 1.57 | 1.90 | 2.26 | 2.80 | 1.12 | 4.37 | 0.23 | 2.66 | 1.10 | Unknown |
| ASD7 | 0.81 | 1.75 | 3.32 | 0.40 | -0.46 | 2.20 | 0.29 | -1.49 | 0.82 | -0.49 | Unknown |
| ASD16 | 1.85 | 1.79 | 0.21 | 0.21 | -0.38 | 0.80 | -0.14 | 3.13 | 1.49 | 0.73 | Unknown |
| ASD20 | -1.07 | 0.27 | 0.09 | -0.03 | 1.58 | 1.51 | -1.38 | -1.00 | -0.62 | 2.80 | Unknown |
| ASD37 | 0.77 | 0.14 | -1.20 | 0.35 | 1.31 | -0.50 | 0.03 | 0.03 | 1.93 | 1.13 | Unknown |

**Figure 3.**
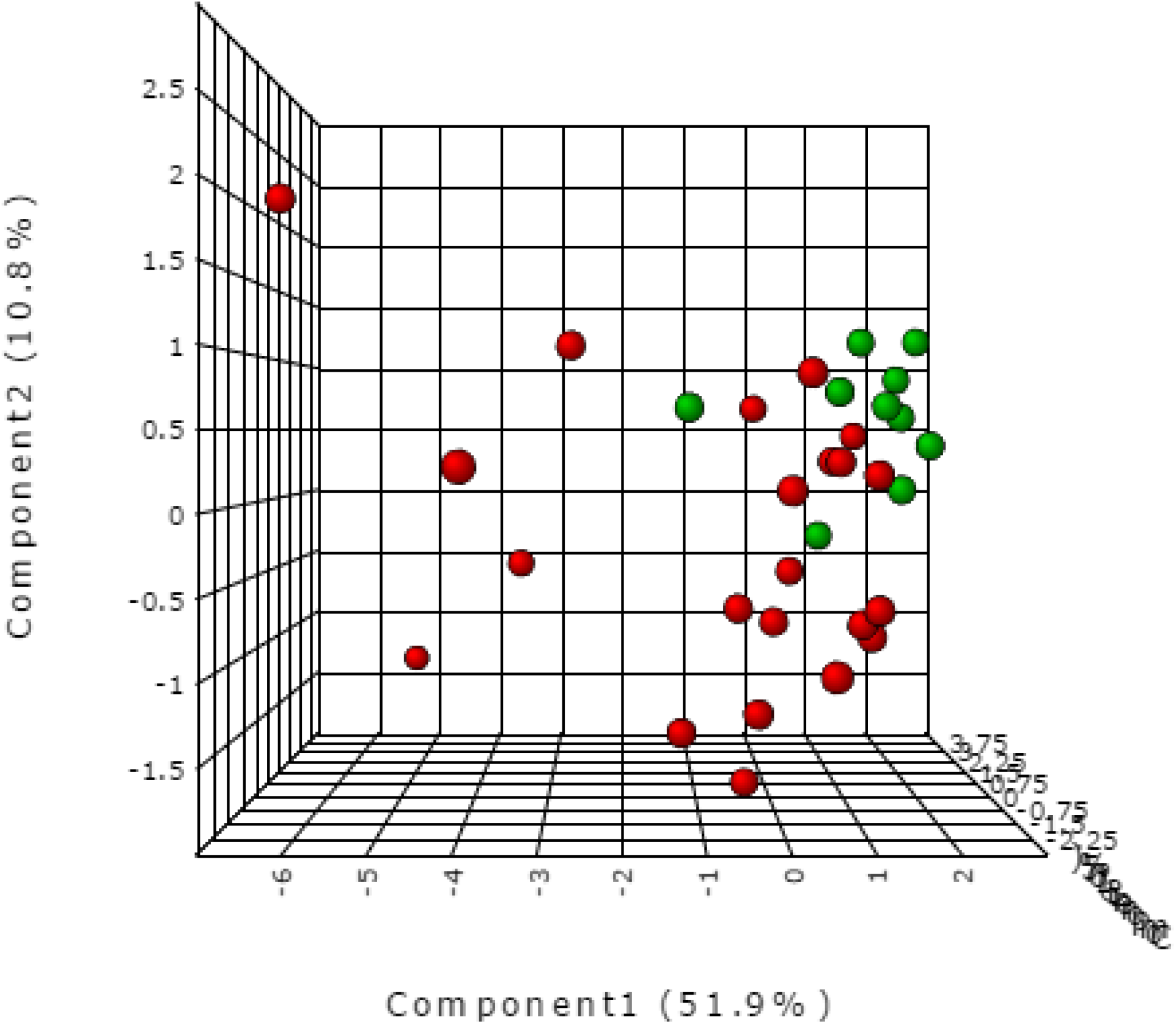
Partial least squares Discriminant Analysis (PLS-DA) of patients with ASD using gene expression data of 10 selected circadian genes. Red balls represent 22 patients with sleep problems reported, and green balls denote 10 patients with normal sleep pattern.

## Discussion

### Roles of circadian genes in the pathogenesis of ASD

We documented that for many patients with ASD, expression levels and gene-to-gene associations of circadian genes in lymphoblastoid cells are different from controls. The molecular changes are likely caused by defects in the circadian system, thus might account for some of the common phenotypes seen in patients with ASD.

The notion that circadian genes may play critical roles in the pathogenesis of ASD is supported by a very recent study of gene polymorphism: Yamagata and co-workers reported that patients with ASD have more mutations that affect gene functions in the circadian-relevant genes than in controls (Yang et al., 2015). Additional supportive evidence came from the research by Hu and co-workers in which they showed severely language-impaired ASD were correlated with multiple differential expressed circadian genes in lymphoblastoid cells (Hu et al., 2009).

Our data suggested that for many patients with ASD, there might exist a link between the sleep problems with abnormal expression levels of circadian genes in lymphoblastoid cells (**Table 2 & Figure 3**). This could simply imply that whatever the common causes of sleep disorders in the neurons are also operating in peripheral tissues such as the lymphoblastoid cells. Alternatively, this could also entail that possible defects in the circadian genes are directly affecting the normal functions of both neurons and peripheral tissues. Regardless the true nature of the causes, our finds suggest that sleep disorders in patients with ASD might be due to defects at the molecular level rather than purely a behavioral presentation.

Tordjman and coworkers proposed that ASD is a disorder of biological and behavioral rhythms, based on their studies on the roles of melatonin in patients with ASD (Tordjman et al., 2015). The hypothesis is intriguing, but it lacks a molecular basis. Two studies looked for genes in the melatonin pathway but failed to establish a strong link between the polymorphisms of melatonin-related genes and ASD (Chaste et al., 2010; Jonsson et al., 2010). On the other hand, our findings of abnormal expressions and abnormal co-expression pattern of circadian genes might just provide a mechanistic underpinning for Tordjman’s hypothesis.

### CLOCK gene and ASD

One of the most noticeable features of the co-expression networks of the ASD cells is the CLOCK gene’s complete loss of connection with other circadian genes. The observation suggests that CLOCK gene and its mediated pathways are operating in a different context in the lymphoblastoid cells of ASD patients. In addition, CLOCK gene is one of most frequently dysregulated circadian genes, as seven out of thirty-eight (18%) in our cohort of patients with ASD showed abnormal ranges of expression for CLOCK genes.

The protein encoded by CLOCK plays a central role in the regulation of circadian rhythms. CLOCK protein also functions as a histone acetyltransferase (Doi et al., 2006). Kino and co-workers demonstrated that CLOCK could bind to human glucocorticoid receptor (hGR) and suppress hGR-induced transcription (Nader et al., 2009). The level of acetylated hGR was also correlated with the expressions of CLOCK-related genes in peripheral blood mononuclear cells (Nader et al., 2009). Taken together, these observations suggest that CLOCK is a negative regulator of action in target tissues (Nicolaides et al., 2014).

The roles of CLOCK gene play in human diseases are highlighted by positive results of polymorphism studies. Different alleles of CLOCK gene were associated with multiple diseases and disorders, including ASD (Yang et al., 2015), Alzheimer’s disease (Chen et al., 2013a; Chen et al., 2013b), ADHD (Jeong et al., 2014), and behavioral changes (Garaulet et al., 2011).

### Individual difference and the implication for personalized medicine

We observed a tremendous variation on the circadian gene expressions in the cells of patients with ASD (**Figure 4**). The variation pointed to both directions of expressions, either dramatically increased or dramatically decreased. For example, among the seven individuals who were outliers for the expression of the CLOCK gene, two of them displayed decreased expressions while five having increased expressions. Similarly, for PER1 gene two patients with ASD showed decreased expressions while five patients having increased expressions. The trend extends to most of the circadian genes under study (**Figure 4**).

**Figure 4.**
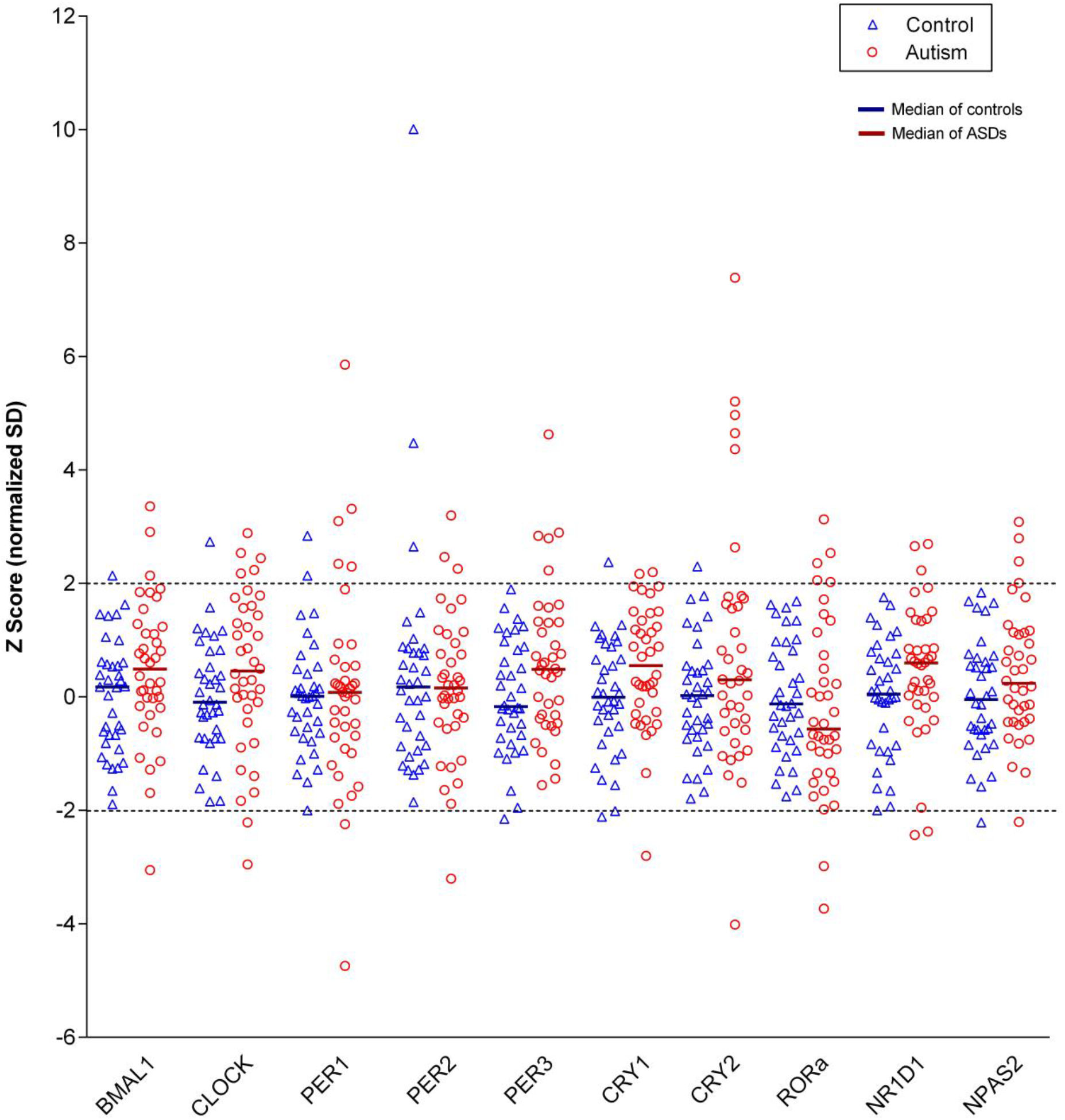
Scatter plot of expression levels of 10 circadian genes in the lymphoblastoid cells of individual controls versus individual patients with ASDs. The dotted lines denote z-scores =± 2.

The individuality and diversity of patients with ASD in expressions of circadian genes we observed here not only reinforce the concept of genetic heterogeneity of ASD, but also necessitate the need for development of personalized medicine for the treatment of ASD.

### Future research and potential for therapeutics

To fully unravel the functional roles of circadian genes in the pathogenesis of ASD, more studies are required. One pressing question perhaps is to ask: what are the causes of the abnormal expression pattern of circadian genes in the cells of patients with ASD? Could it be some genetic mutations in the circadian genes themselves? Are there defects in other regulatory genes in the circadian pathway yet to be discovered?

Another important question will be the biological consequences of dysregulated circadian genes in the cells of patients with ASD. Are metabolisms of bioenergetic molecules such as sugar or lipid disturbed in cells of the patients with ASD? Do the cytokine profiles in cells of the patients with ASD change?

Our study utilized a design in which the case-control cohort was matched with gender, ethnicity, and age (see **Materials and Methods** for details). To deepen the understanding of the molecular mechanisms of circadian genes and circadian rhythm in ASD, future studies should extend to a large cohort with diverse background.

## Materials and Methods

### Sample selection and cell line culture

Thirty-eight patients with non-syndromal ASDs and thirty-six controls were selected for the study. Because ASD is a heterogeneous disease, in order to reduce the confounding effect of various non-genetic factors and to increase the signal/noise ratio of the assays, the case-control cohort was matched with age, ethnicity, and gender: 2-20 year old, Caucasian, and males. The age distribution can be found in **Supplementary Figure 1**. The non-syndromal patients were part of the South Carolina Autism Project (SCAP) and were diagnosed with autistic disorder based on evaluation using the Autism Diagnostic Interview-Revised (ADI-R) and using the American Psychiatric Association’s diagnostic and statistical manual of mental disorders-fourth edition-text revision (DSM-IV-TR) (American Psychiatric Association, 2000). Informed consent was approved by the Self Regional Healthcare Institutional Review Board (IRB) for Human Research and was reviewed and signed by all the participants evaluated and/or their legal guardian.

Genetic tests excluded major chromosomal abnormalities, Fragile X syndrome, Rett syndrome (by testing the MECP2 gene), and abnormalities in plasma amino acid levels.

Cell lines were obtained by immortalization of lymphocytes from blood samples using Epstein-Barr virus. The lymphoblastoid cell lines were maintained until harvesting in RPMI-1640 medium (Corning, Corning, NY, USA), supplemented with 15% fetal bovine serum (Atlanta Biological, Lawrenceville, GA, USA), 2mM L-Glutamine, 1% penicillin/streptomycin and 1% MEM Non-essential Amino Acid Solution (all three reagents were purchased from Sigma-Aldrich, St. Louis, MO, USA). Cells were cultured at a density lower than 1 × 10^6^/ml to ensure logarithmic growth.

### Real-Time Quantitative PCR (qPCR)

Total RNA was isolated using the RNeasy Mini RNA isolation kit (Qiagen, Valencia, CA) according to the manufacturer’s instructions. Reverse transcription was performed using Verso cDNA kit (Thermo Fisher Scientific, Waltham, MA, USA). Real-time quantitative PCR (qPCR) for quantification of mRNA was conducted using iQ SYBR Green Supermix and an BioRad CFX96 Real-time PCR System (BioRad, Hercules, CA). The gene-specific primers and their cycler conditions used for qPCR were adopted from literatures or designed using the Primer3 online tool (Rozen and Skaletsky, 1998). The information for primer sequences was listed in **Supplementary Table 1**. The primer DNA oligos were purchased from Eurofins MWG Operon (Huntsville, AL, USA). Our preliminary analysis suggested that mRNAs for the glyderaldehyde-3-phosphate dehydrogenase (GAPDH) gene do not vary among the control samples. The GAPDH gene was therefore used as the reference gene. Genes of interest and reference genes (i.e., GAPDH) were PCR amplified for construction of standard curves. Each sample was analyzed in triplicate.

### Data processing and statistical analysis

We used standard curves method to calculate the relative copy numbers of the genes of interest and reference genes (GAPDH) in the qPCR experiments. After normalized to the reference gene, the expression level of each individual gene was presented as ratio to the average of controls. Z-scores were computed using the means and standard deviations of the control samples.

The odds ratio was calculated using the statistics package Prism 6 (GraphPad Software, La Jolla, CA). Pair-wise Spearman’s rank correlation coefficients were calculated for the log2-transformed data using an R script. Co-expression networks between genes were constructed using gene pairs with absolute correlation value equal to larger than 0.6 (*P* value <0.001). The network was visualized using the software CytoScape V3.2.1 (Shannon et al., 2003). For the analysis of the correlation of circadian genes with sleep pattern of patients with ASD, we took advantage of the Z score being a normalized measurement and treated the absolute value of a Z-score as a distance from the mean. Partial least squares Discriminant Analysis (PLS-DA) was then conducted using the on-line software MetaboAnalyst 3.0 (Xia et al., 2015).

## Acknowledgements

The experimental work was performed while the authors were at the J.C. Self Research Institute of Human Genetics, The Greenwood Genetic Center, Greenwood, South Carolina. We are grateful for the support provided by the Greenwood Genetic Center. We thank Cindy Skinner who was the research coordinator at Greenwood Genetic Center for organizing patients’ information.

## Supplementary Tables and Figures

**Supplementary Table 1.**
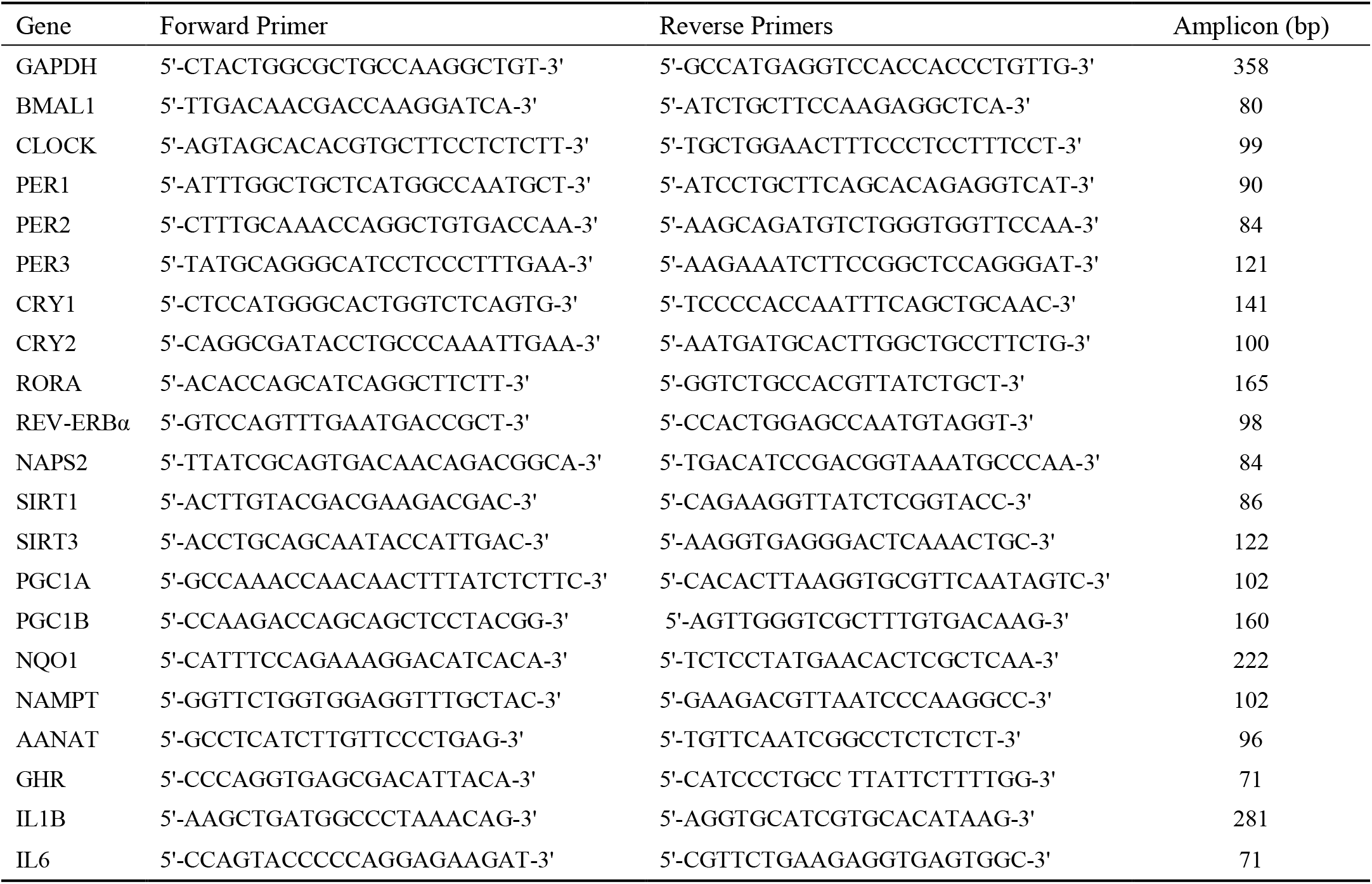
qPCR primer sequences and amplicon sizes.

**Supplementary Figure 1.**
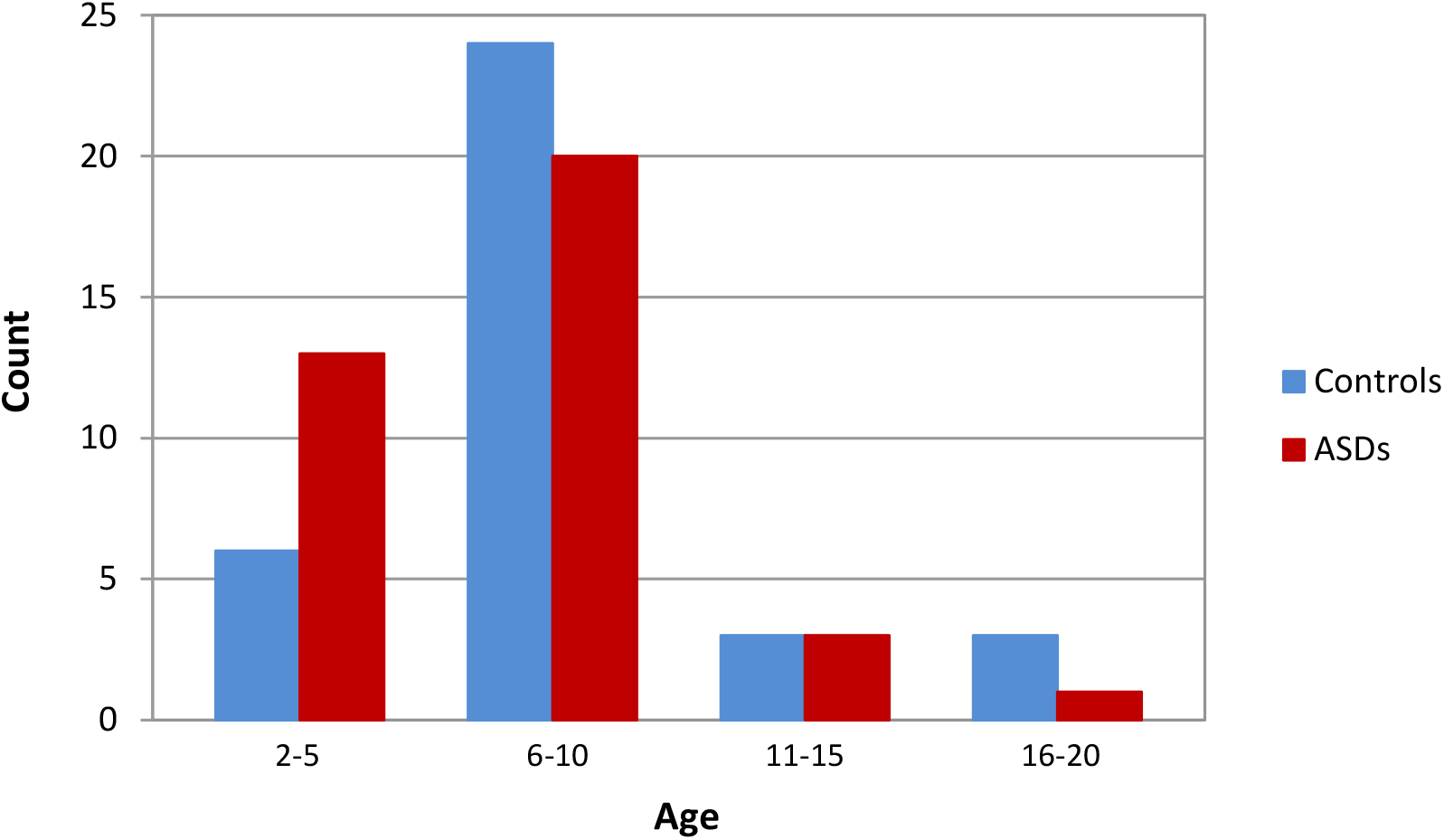
Age distribution of the study cohort.

## Notes

### Competing Interest Statement

The authors have declared no competing interest.

